# Combinations of Peptides Synergistically Activate the Regenerative Capacity of Skin Cells In Vitro

**DOI:** 10.1101/2021.07.07.451479

**Authors:** Michael J. Flagler, Makio Tamura, Tim Laughlin, Scott Hartman, Julie Ashe, Rachel Adams, Kim Kozak, Kellen Cresswell, Lisa Mullins, Bradley B. Jarrold, Robert J. Isfort, Joseph D. Sherrill

## Abstract

**OBJECTIVE:** To explore synergistic effects related to skin regeneration, peptides with distinct biological mechanisms of action were evaluated in combination in different skin cell lines in the presence or absence of niacinamide (Nam). Furthermore, the synergistic responses of peptide combinations on global gene expression were compared to the changes that occur with fractional laser resurfacing treatment, a gold standard approach for skin rejuvenation, to further define optimal peptide combinations.

**METHODS:** Microarray profiling was used to characterize the biological responses of peptide combinations (+/− Nam) relative to the individual components in epidermal keratinocyte and dermal fibroblast cell lines. Cellular functional assays were utilized to confirm the synergistic effects of peptide combinations. Bioinformatics approaches were used to link the synergistic effects of peptide combinations on gene expression to the transcriptomics of the skin rejuvenation response from fractional laser treatment.

**RESULTS:** Microarray analysis of skin cells treated with peptide combinations revealed synergistic changes in gene expression compared to individual peptide controls. Bioinformatic analysis of synergy genes in keratinocytes revealed activation of NRF2-mediated oxidative stress responses by a combination of Ac-PPYL, Pal-KTTKS, and Nam. Additional analysis revealed direct downstream transcriptional targets of NRF2/ARE exhibiting synergistic regulation by this combination of materials, which was corroborated by a cellular reporter assay. NRF2-mediated oxidative stress response pathways were also found to be activated in the transcriptomics of the early skin rejuvenation response to fractional laser treatment, suggesting the importance of this biology in the early stages of tissue repair. Additionally, a second combination of peptides (pal-KT and Ac-PPYL) was found to synergistically restore cellular ATP levels that had been depleted due to the presence of ROS, indicating an additional mechanism whereby peptide synergies may accelerate skin repair.

**CONCLUSION:** Through combinatorial synergy studies, we have identified additional in vitro skin repair mechanisms beyond the previously described functions of individual peptides and correlated these to the transcriptomics of the skin rejuvenation response of fractional laser treatment. These findings suggest that specific peptides can act together, via complementary and synergistic mechanisms, to holistically enhance the regenerative capacity of in vitro skin cells.

## INTRODUCTION

Both intrinsic and extrinsic skin aging are characterized by a progressive deterioration of skin’s youthful structure (1). Aging-induced structural changes occur across all compartments of the skin, from deep within the dermis, to the superficial viable epidermis and stratum corneum layers (1, 2). Hallmark structural changes include a loss and breakdown of dermal collagen, elastosis, flattening of rete ridges at the dermal-epidermal interface, and thinning of the viable epidermis (1, 3–5). Therefore, a holistic solution for repairing the structural damage associated with skin aging must act on both epidermal and dermal layers to address these diverse structural changes.

Many cosmetic procedures have been developed to rejuvenate aged skin by activating the skin’s innate wound repair processes in both the epidermis and dermis. Such modalities exhibit diverse modes of action, ranging from chemical damage, to physical damage, to thermolysis, with the common denominator being that these approaches all inflict damage to the skin to trigger activation of a wound healing response (6–13). Recently, thermal energy-based approaches (i.e., radiofrequency (e.g., Thermage™), ultrasound (Ultherapy™), and lasers (e.g., Fraxel™)), which act via the same general mode of action on a continuum, based on energy and time duration, have become popular options. Laser-based approaches are now widely considered a gold standard treatment for aging related skin rejuvenation (6, 7). These approaches, however, come with the trade-off that the skin must first be damaged to achieve the resultant rejuvenation and appearance benefits. Such procedures can lead to long healing times and significant side effects, creating a need for alternative approaches which activate the biological healing responses in skin without the need to inflict injury to the tissue.

Topically applied bioactive materials have been shown to activate various repair processes associated with dermal and epidermal wound healing, with biologically active peptides amongst the most prevalent class of such materials. Collagen-derived peptides such as Pal-KTTKS (Promatrixyl™) and Pal-KT (Palestrina™), for instance, have been shown to stimulate new collagen synthesis in vitro (14–16). These peptides have been proposed to act as signaling agents to fibroblasts, mimicking damage to collagen such as occurs during wounding which can cause collagen fragmentation into peptides (15). Thus, these peptides (often referred to as matrikines) offer one approach to stimulate in vitro collagen synthesis without actual wounding having taken place (17). Carrier peptides represent another general category of peptides which stabilize and deliver trace elements to cells including copper and manganese, which are necessary for enzymatic processes involved in tissue repair and wound healing (18, 19). GHK-Cu, for example, is reported to facilitate the delivery of copper ions to target enzymes that use copper as a co-factor, including enzymes (e.g., lysyl oxidase) involved in dermal matrix stabilization (20–22). Finally, while these peptides activate repair processes in the dermis, another peptide, Ac-PPYL (Syniorage™) has been reported to stimulate in vitro biological processes associated with epidermal repair (23). Specifically, this 4-amino acid fragment of PR-39, a proline-rich antimicrobial peptide, has been shown in vitro to induce keratinocyte cell growth and promote epidermal cohesion via syndecan-1 and collagen XVII synthesis (24–27).

We hypothesized that combining skin repair peptides to target both epidermal and dermal repair processes, alone or in combination with the nicotinamide adenine dinucleotide (NAD+) precursor niacinamide (Nam), would lead to more holistic in vitro rejuvenation through working across multiple skin layers. Additionally, we explored the possibility that combinations of such peptides may lead to new, previously uncharacterized in vitro biological responses associated with skin repair, and/oract in a synergistic mannerto deliver enhanced benefits overthe individualpeptide constituents. In the present study, we identified two such combinations of materials: Ac-PPYL + Pal-KTTKS + Nam, and Ac-PPYL + Pal-KT, which act via both complementary mechanisms as well as synergistically activating in vitro pathways associated with the early skin rejuvenation response to fractional laser treatment. These complexes may serve as future optimized peptide combinations to achieve enhanced anti-aging benefits.

## MATERIALS & METHODS

### Microarray Profiling

Microarray profiling was performed as previously described (28). In brief, tert-immortalized keratinocytes (kindly provided by Jerry Shay, UT-Southwestern) or immortalized dermal fibroblasts (BJ) (ATCC, CRL-2522™) were treated with niacinamide (500μM), Ac-PPYL (0.5 μg/mL), Pal-KTTKS (1.0 μg/mL), or the tri-combination for 6 hr (n = 5 per treatment group). Cells were then lysed in RLT Buffer (QIAGEN, Hilden, Germany) and total RNA was extracted using the Agencourt RNAdvance Tissue kit (Beckman Coulter, A32646). Biotinylated cRNA libraries were synthesized from 250 ng of total RNA using the Affymetrix HT 3’ IVT Plus kit (Affymetrix, Santa Clara, CA) and the Beckman Biomek FXp Laboratory Automation Workstation (Beckman Coulter, Brea, CA). Biotinylated cRNA was subsequently fragmented by limited alkaline hydrolysis and then hybridized overnight to the Affymetrix Human Genome U219-96 GeneTitan array. Plates were scanned using the Affymetrix GeneTitan Instrument utilizing AGCC GeneTitan Instrument Control software version 4.2.0.1566.

The clinical design for laser resurfacing using Fraxel^®^ was approved by the Schulman Associates Institutional Review Board.14 women aged 30-55 with Fitzpatrick skin type I-IV underwent fractional laser treatments on the back using a Fraxel^®^ laser. Full thickness skin punch biopsies were obtained at baseline (untreated, day 0) and 3 days post-treatment. Microarray profiling was performed using the HG-U219 array (Affymetrix) as similarly described (3). Flash-frozen, bisected 4mm full-thickness skin punch biopsies were homogenized in TRIzol reagent (ThermoFisher, Waltham, MA) and total RNA was extracted according to the manufacturer’s protocol. RNA was further purified using RNEasy spin columns (QIAGEN, Hilden, Germany). Biotinylated cRNA libraries were synthesized from 500 ng of total RNA using the Affymetrix HT 3’ IVT Express kit (Affymetrix, Santa Clara, CA) and ta Beckman Biomek FXp Laboratory Automation Workstation (Beckman Coulter, Brea, CA). Biotinylated cRNA was subsequently fragmented by limited alkaline hydrolysis and then hybridized overnight to the Affymetrix Human Genome U219-96 GeneTitan array. Plates were scanned using the Affymetrix GeneTitan Instrument utilizing AGCC GeneTitan Instrument Control software version 4.2.0.1566. Image data was summarized using Affymetrix PLIER algorithm with quantile normalization.

### Bioinformatic Analyses

U219 microarray chip data from the in vitro keratinocyte study is normalized by the Single-channel array normalization (SCAN) and Universal exPression Codes (UPC) method using the Brainarry lab’s custom CDF (29, 30). In this way, it is possible to have a clear one-to-one mapping between probe sets and gene and to enable comparing data across different batches by removing the probe- and array-specific background noise. The log2 fold change, and its 95% confidence intervals and p-value of each treatment compared to the control were computed by using the Limma package in Bioconductor (31). A gene showing a potential synergistic effect was evaluated by several criteria; 1) the peptide combination exhibited a significant difference (p-value < 0.05) compared to the control, 2) the lower or upper confidence interval of the log2 fold change of the combination treatment was larger or smaller, respectively, than the sum of each treatment effect with respecting the directionality of changes, and 3) the expression level of the gene was more than the lower 30% percentile among those of all genes to confirm that the gene was truly expressed.

Microarray data from untreated and laser resurfacing-treated skin underwent rigorous quality check using methods and metrics described by Canales, et al, and data were fitted to a linear mixed ANOVA model using SAS software PROC GLIMMIX, version 9.4 of the SAS System for Windows (32). Gene expression (in log scale) of the 3 days post-treatment site was compared to that of the baseline (day 0) untreated site. Subject, Affymetrix plate (samples from 14 subjects were processed on two plates with seven subjects per plate) and Site were class variables. Subject (nested within Plate) was the random effect in the model.

Gene signature analyses were performed using GeneSpring version 14.8 (Agilent Technologies, Santa Clara, CA). Hierarchical clustering of the log2 fold change values using Euclidean distance was performed to create the dendrograms. Biological pathway enrichment was performed on the synergy gene signature using Ingenuity Pathway Analysis version 01-07 (QIAGEN). Fisher’s exact t-test was used to assess significantly associated canonical signaling pathways, and z-scores were calculated to predict pathway activation states (activated pathways, z-score >0; inhibited pathways z-score <0).

### NRF2 Reporter Assay

The ARE32 reporter cell line was generated in Roland Wolf’s laboratory and obtained from CXR BioSciences (33). The cells were grown in maintenance medium (DMEM, Gibco 11054-020) supplemented with 0.8 mg/ml gentamicin (Gibco 10131-027), 1:100 dilution of GlutaMax (Gibco 15140-148), 10% HI FBS (Gibco A31604), and pen/strep and seeded into 96-well plates 24 hr prior to treatment. Media was then replaced with fresh treatment media (phenol red free DMEM, Gibco 11054-020) with 1:100 dilution of GlutaMax and penicillin/streptomycin and cells were treated (n = 3 per group) with compounds and tBHQ (Sigma 11294-1) for 24 hr. Post treatment, cells were rinsed with 1X PBS, lysed and luminescence was measured according to the manufacturer’s protocol (Promega) using a SynergyNeo plate reader (Biotek). Treatment values are reported as fold induction over vehicle control. Paired group comparisons were made using the T-test function, two tail with even distribution functions in Excel. P-values are adjusted by the Bonferroni correction of multiple hypothesis testing, and the adjusted p-values ≤ 0.05 were considered statistically significant.

### Keratinocyte ATP under Peroxide Stress Assay

Tert-immortalized keratinocytes (kindly provided by Jerry Shay, UT-Southwestern) were cultured in EpiLife Medium (catalogue # MEPICFPRF500, Thermo Fisher Scientific) supplemented with human keratinocyte growth supplement (catalogue #S-001-5, Thermo Fisher Scientific) and 1% penicillin/streptomycin (catalogue #15140-122, Thermo Fisher Scientific). Cells were split into 96-well plates and treated as follows (n = 3 per group): (i) no stress and no treatment; (ii) stress only; and (iii) stress and treatment. For the stressed samples (groups (ii) and (iii)), culture medium containing 400 μM hydrogen peroxide (H_2_O_2_) was used. For the stress and treatment samples (group (iii)), treatments were applied immediately after H_2_O_2_. Stressed/treated cells were then returned to a CO_2_ incubator for 60 minutes. ATP levels were then measured using the ATP Glo^®^ reagent (Promega, Madison WI) and luminescence recorded using an ENVISION plate reader (Perkin Elmer) according to the manufacturer’s protocol. Paired group comparisons were made using the T-test function, two tail with even distribution functions in Excel. P-values ≤ 0.05 were considered statistically significant. For cases in which two treatments appeared to synergistically enhance ATP levels, the results of the combined treatments were compared to the sum of the individual treatments to determine a synergy factor. The synergy factor was calculated as the ratio of the observed effect from the combined treatments divided by the sum of the results from each individual treatment. A synergy factor of 1.00 indicates that the combination treatment performed as expected whereas synergy factors > 1.00 indicate a synergistic benefit beyond simple additivity/expectation.

### QPCR Analysis

QPCR was used to validate the synergistic action of the niacinamide + Ac-PPYL + Pal-KTTKS combination or Ac-PPYL + Pal-KT combination on a subset of the transcripts identified from the microarray profiling. Immortalized keratinocytes (tKC) or fibroblasts (BJ) were treated for 24 hr then the cells were lysed in RLT buffer(Qiagen) and total RNA purified using the Biomek FxP and the RNAdvance 2.0 Tissue Isolation kit (Beckman Coulter, A32646). cDNA was generated using 500 ng of total RNA and the High-Capacity cDNA with Reverse Transcription kit (Applied Biosystems p/n 4368814). QPCR was performed using the Wafergen SmartChip system (TakaraBio, p/n 640036). Data were normalized to the arithmetic mean of five housekeeping genes (ACTB, B2M, GAPDH, GUSB and PPIA). A linear model was fit comparing treatments using the limma R (Version 4.0.4) package. For each treatment, a total of n=6 replicates were used. Adjusted fold-change values between treatments were extracted from the model and the empirical Bayes method was used to extract p-values (2-sided equal variance) for significance testing.

## RESULTS

In an effort to identify synergistic peptide combinations with the potential to stimulate skin repair processes, we performed microarray profiling to characterize the genome-wide transcriptional response of keratinocytes in vitro to a peptide combination of Ac-PPYL and Pal-KTTKS with Nam relative to the individual components. We identified a gene expression signature composed of 857 genes that exhibited synergistic activation in the presence of Ac-PPYL, Pal-KTTKS, and Nam (Fig 1A, B). Candidate synergy genes displayed a diverse array of biological functions with direct linkages to repair processes. TIMP1, for instance, has been reported to play a direct role in protection and recovery from both dermal and epidermal photodamage due to its suppression of extracellular matrix (ECM) degradation and inflammation (34).

**Figure 1.**
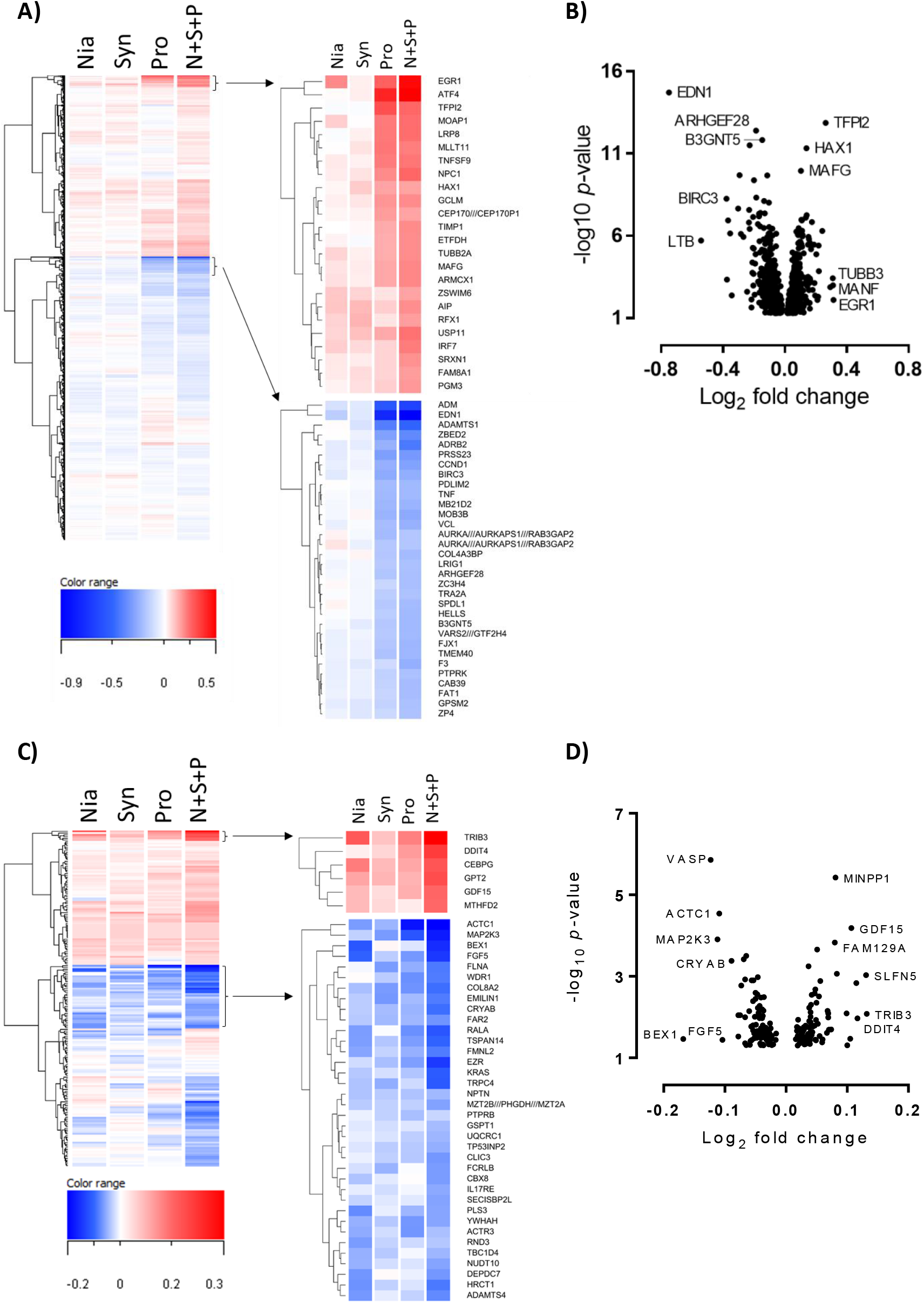
Synergistic effects of niacinamide, Ac-PPYL, and Pal-KTTKS on gene expression in keratinocytes and fibroblasts. **A)** Dendrogram displaying hierarchical clustering of genes by average log_2_ fold change (compared to untreated keratinocytes at 6hr) that showed synergism in the niacinamide (Nia), Ac-PPYL (Syn), and Pal-KTTKS (Pro) tri-combination treatment (N+S+P) (n = 857 total). Clusters of the top upregulated (upperright) and downregulated (lowerright) genes are shown. **B)** Volcano plot showing –log10 *p*-value and log2 fold change for N+S+P treated keratinocytes (compared to untreated keratinocytes at 6hr). Labelled are the top 3 most significant and top 3 fold changes (up- and downregulated) synergy genes. **C)** Dendrogram displaying hierarchical clustering of genes by average log2 fold change (compared to untreated fibroblasts at 6hr) that showed synergism in the niacinamide (Nia), Ac-PPYL (Syn), and Pal-KTTKS (Pro) tri-combination treatment (N+S+P) (n = 179 total). Clusters of the top upregulated (upper right) and downregulated (lower right) genes are shown. D) Volcano plot showing –log10 *p*-value and log2 fold change for N+S+P treated fibroblasts (compared to untreated fibroblasts at 6hr). Labelled are the top 3 most significant and top 3 fold changes (up- and downregulated) synergy genes.

To further investigate the in vitro biological processes synergistically activated by the Ac-PPYL, Pal-KTTKS, and Nam combination, we assessed canonical signaling pathways for biological enrichment in the synergistic gene signature. Interestingly, the NRF2-mediated oxidative stress response was found to be one of the most strongly activated synergy pathways by this combination of materials (p = 2.58 × 10^−4^, z-score = 2.53) (Fig 2A). Nuclear factor E2-related factor 2 (NRF2) has been identified as one of the primary transcription factors which acts on a cis-acting DNA element termed the antioxidant response element (ARE) (35). ARE sequences are found in the promoters of many genes whose protein products are involved in the detoxication and elimination of reactive oxidants and electrophilic agents, as well as enhancing cellular antioxidant capacity, making activation of the NRF2-ARE signaling pathway a key cellular defense mechanism against oxidative stress-induced damage (36, 37). A focused analysis of NRF2 target genes revealed many direct downstream transcriptional targets of NRF2/ARE which exhibit synergistic regulation by this combination of materials (Fig 2B), encompassing a spectrum of in vitro cellular repair processes ranging from damage neutralization and mitigation to renewal processes. For example, NRF2 target genes involved in immediate reactive oxygen species (ROS) neutralization (*SOD1*, *GPX2*, *NQO2*), glutathione synthesis (*GCLM*), and protein damage repair via the proteasome (*VCP*) were found to be synergistically regulated by this combination of materials (Fig 2B).

**Figure 2.**
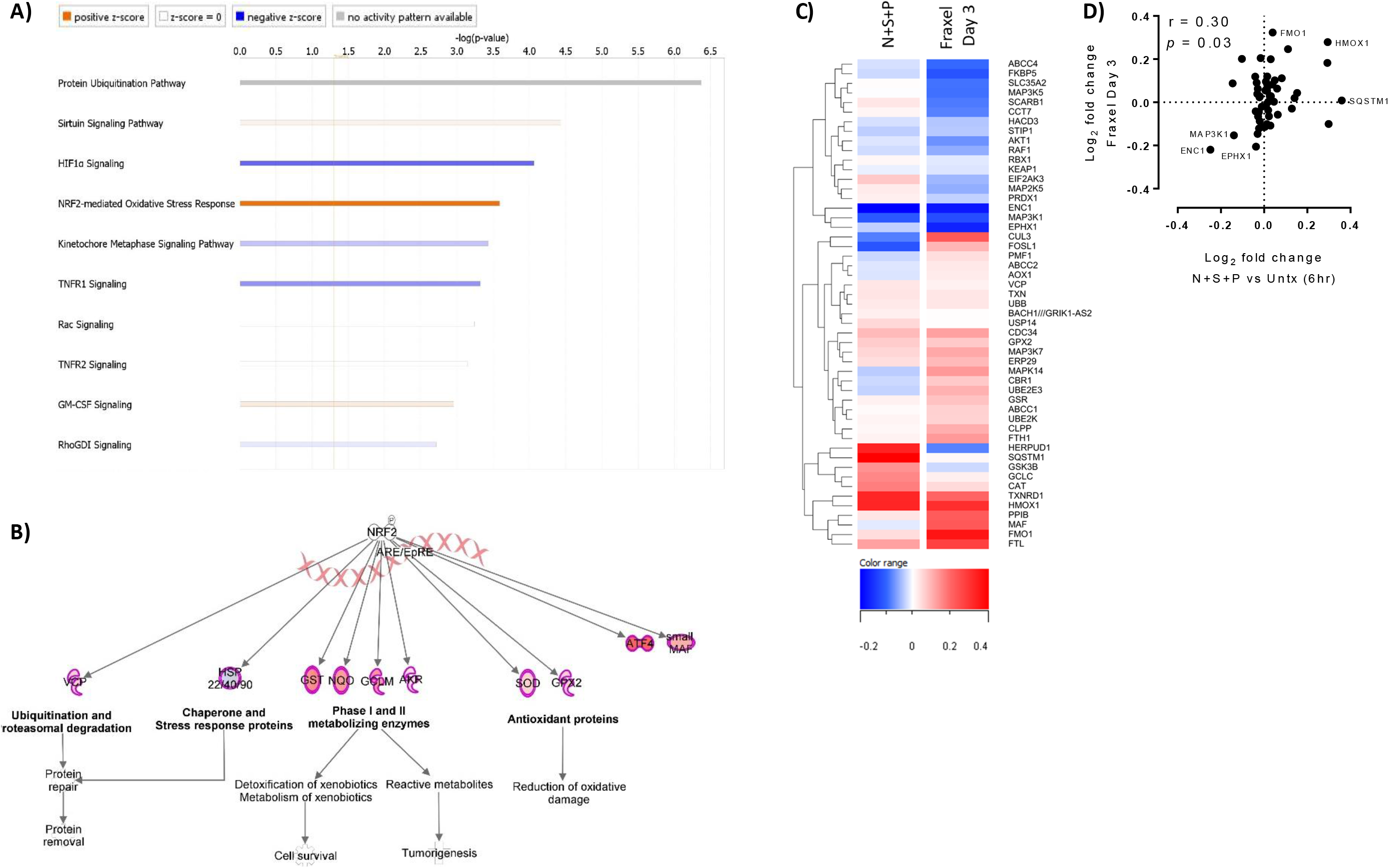
Activated oxidative stress response links synergy genes and transcriptomics of early skin rejuvenation response. **(A)** Top 10 canonical signaling pathways associated with the synergy genes in tert-KCs (by Fisher’s –log_10_ *p*-value). Bar coloration reflects the degree to which pathways are predicted to be activated (increasing orange = increasing z-score), inhibited (increasing blue = decreasing z-score), unchanged (white, z-score = 0), or, for pathways for which no activity prediction can be made (gray, z-score = n/a). **(B)** Direct downstream transcriptional targets of NRF2 binding to the ARE in DNA (pink Xs) that show synergistic regulation by niacinamide (Nia), Ac-PPYL (Syn), and Pal-KTTKS (Pro) (N+S+P) (red, upregulated; blue, downregulated)**. (C)** Dendrogram displaying hierarchical clustering of all genes within the NRF2-mediated oxidative stress responses canonical pathway (n = 53) by average log_2_ fold change in the synergy genes (N+S+P, compared to untreated keratinocytes at 6hr) and in skin 3 days post-laser resurfacing treatment (Fraxel Day 3, compared to baseline untreated skin at Day 0). **(D)** Log_2_ fold change plot and Pearson correlation for all 53 NRF2 genes from (C) between the synergy treatment (N+S+P, compared to untreated keratinocytes at 6hr) and in skin 3 days post-laser resurfacing treatment (Fraxel Day 3, compared to baseline untreated skin at Day 0).

To explore how the NRF2 response activation profile induced by the Ac-PPYL, Pal-KTTKS, and Nam combination compared to that of rejuvenated skin, we compared the in vitro modulation of all genes in the NRF2 canonical signaling pathway (n=53) by the material combination to the in vivo response of skin post-laser resurfacing treatment. The combination elicited a similar response profile as observed in skin 3 days post-laser resurfacing treatment (compared to baseline) (Fig 2C), with a statistically significant positive correlation observed (p= 0.03) (Fig 2D). These findings suggest both the importance of the NRF2 pathway in the early-stage skin rejuvenation response following a benchmark dermatological procedure, and the potential ability of a combination of bioactive materials to elicit a similar response without inflicting a wound.

To validate the NRF2 activation by the bioactive combination, a functional assay based on a reporter system was employed. This system consists of MCF7 cells transfected with the promoter region of GST (an NRF2 target) upstream of the luciferase coding sequence. In this assay, activation of NRF2 results in increased production of luciferase, which is detected as luminescence output. Synergistic in vitro activation of an NRF2 target gene promoter by the combination of Ac-PPYL, Pal-KTTKS, and Nam was observed in this reporter system, with higher luminescence observed for the combination than the expected additive response of the individual components. Interestingly, this synergy was observed to be further augmented by the incorporation of two additional peptides (Pal-KT and RGS) into the complex and could be reproduced by the combination of four peptides (Pal-KTTKS + Ac-PPYL + Pal-KT + RGS) alone in the absence of Nam (Fig 3).

**Figure 3.**
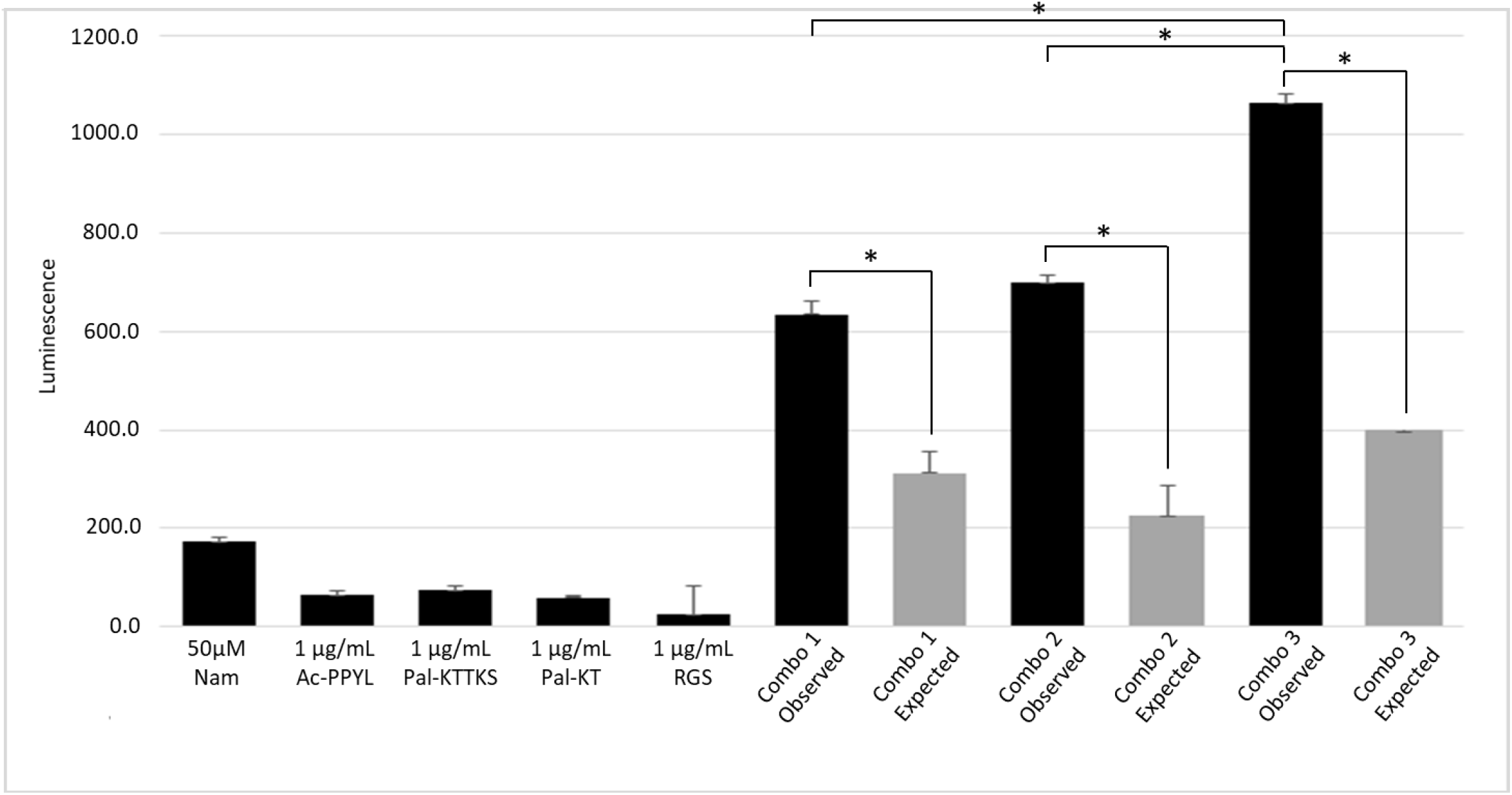
Synergistic activation of *NRF2* target gene promoter by peptide combinations. ARE32 *NRF2* reporter cells were treated with Nam (50 μM), Ac-PPYL(1 μg/mL), Pal-KTTKS (1 μg/mL), Pal-KT (1 μg/mL), and RGS (1 μg/mL), alone or in combination. ARE activation (i.e., luminescence) was measured 24 hr post-treatment (*, adjusted p-value < 0.05). Combo 1 refers to Nam + Pal-KTTKS + Ac-PPYL; Combo 2 refers to Pal-KTTKS + Ac-PPYL + Pal-KT + RGS; Combo 3 refers to Nam + Pal-KTTKS + Ac-PPYL + Pal-KT + RGS.

In addition to synergistic NRF2 activation in keratinocytes, the biological synergy of the Ac-PPYL, Pal-KTTKS, and Nam combination was also identified in dermal fibroblasts. Using the same microarray profiling approach as performed in keratinocytes, an additional 179 transcripts were found to be synergistically regulated in fibroblasts (Fig 1C, D), four of which were subsequently validated by qPCR (Supplemental Table I). Notably, these transcripts included *SLFN5*, which dampens MMP expression and release to initiate the ECM rebuilding phase, and *GDE1*, which codes for an enzyme that produces fatty acid amides which dampen inflammatory processes (38, 39).

A second combination of peptides targeting both epidermal and dermal repair processes, Ac-PPYL and Pal-KT, was also evaluated for in vitro activation of beneficial biological functions beyond those of the individual constituents. In addition to the previously described roles of these peptides in skin repair, additional synergy candidate genes were identified via microarray profiling in dermal fibroblasts and validated by qPCR (Supplementary Table II). Amongst the genes synergistically activated by this peptide combination were genes involved in lipid metabolism and energy homeostasis. Notably, this peptide combination was shown to upregulate *PPARA*, which codes for the nuclear receptor protein peroxisome proliferator activated receptor alpha (PPAR-alpha), a transcription factor and a major regulator of lipid metabolism and glucose homeostasis (40–42). As energy is required to fuel the repair and rebuilding processes involved in cellular regeneration, we further explored the ability of this peptide combination to improve cellular ATP production in cells subjected to oxidative stress. The combination was found to synergistically restore in vitro cellular ATP levels that had been depleted due to the presence of ROS (Fig 4), providing additional functional support for the synergistic activation of cellular energy associated pathways in skin cells.

**Figure 4.**
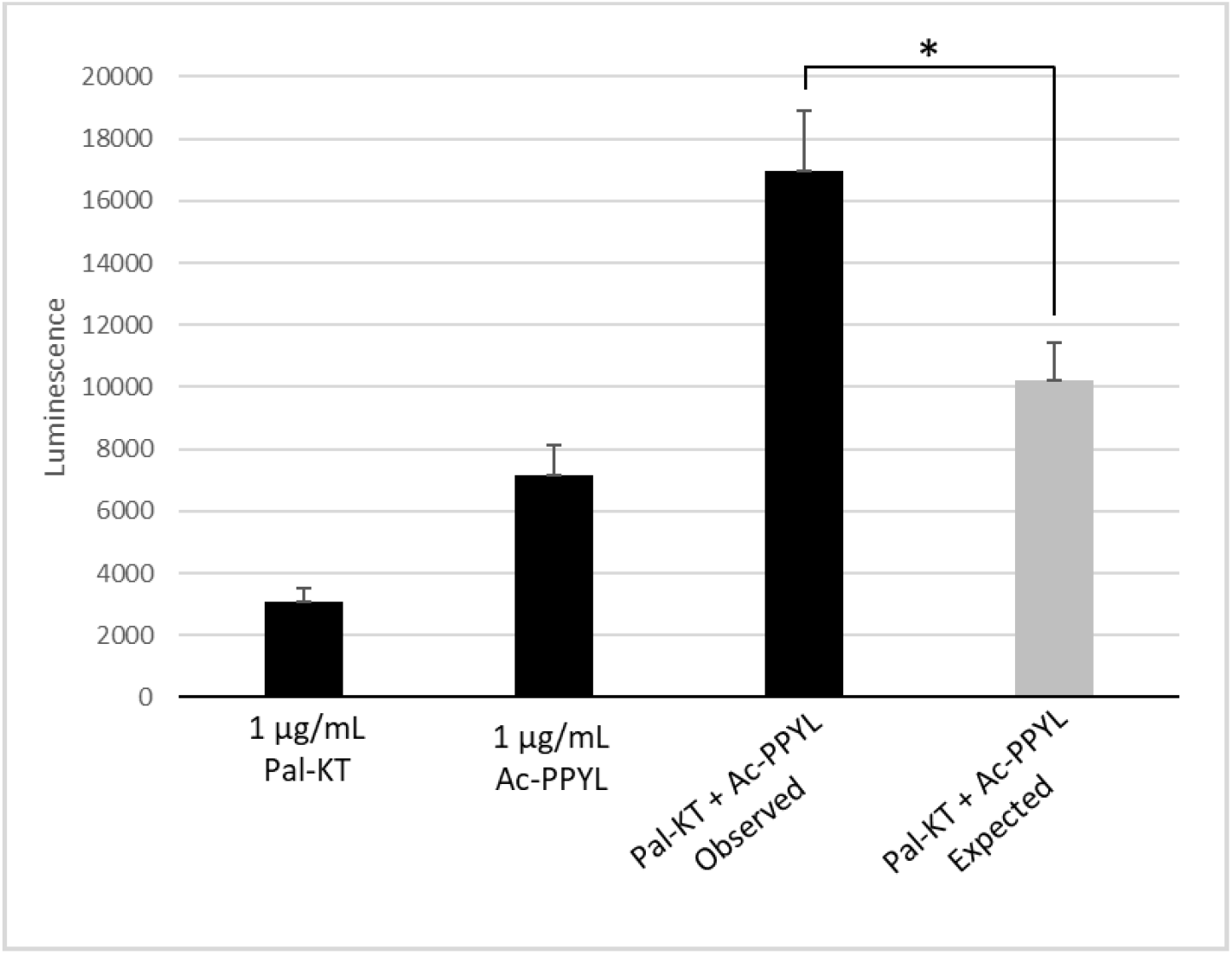
Ac-PPYL and Pal-KT peptides synergistically stimulate ATP levels in stressed keratinocytes. H_2_O_2_-stressed tert-KCs were treated with Ac-PPYL (1 μg/mL), Pal-KT (1 μg/mL), and Ac-PPYL + Pal-KT combination for 1 hr prior to ATP measurement (*, p-value < 0.05).

## DISCUSSION

Skin is the first line of defense against environmental insults that would otherwise damage sensitive underlying tissue and organs. As such, skin cells tend to be more at-risk from oxidative stress than many other cell types in the body due to their high exposure to ultraviolet radiation and other exogenous stressors. Skin is made up of a variety of different cells that function together in a dynamic, complex manner to maintain the health and homeostasis of the organ. However, as skin cells age or become damaged, they can lose their ability to function at the level needed to maintain young, healthy-looking skin (3, 43). The tell-tale signs of skin aging, such as fine lines, wrinkles, and sagging skin, are a consequence of the progressive accumulation of damage in skin cells over time (44). As a result, cosmetic rejuvenation procedures have been developed for treating the signs of aging in skin (e.g., Thermage™, Ultherapy™, and Fraxel™), many of which act via inflicting a wounding stimulus to activate the skin’s innate repair machinery. Such procedures, while yielding gold standard anti-aging appearance benefits, are associated with many negative side effects and can require long recovery times (45). Therefore, it would be advantageous to identify skin rejuvenation technologies that could potentially mimic the beneficial effects of these approaches, without having to inflict additional damage to the already compromised skin structure.

Skin cells can be damaged by a variety of endogenous and exogenous factors, including ultraviolet radiation, pollution, and hormonal alterations associated with psychological stress (46). These stressors can cause the production of ROS, which both interfere with normal cellular processes as well as directly damage the molecular building blocks of the cell (DNA, proteins, and lipids) (43). In response, cells have evolved defenses to combat ROS; however, the cell’s natural defenses can be overwhelmed by elevations in stressor-induced ROS, leading to both acute and chronic alterations in cellular homeostasis (47). As ROS accumulate over time, they contribute to oxidative stress at the cellular level, ultimately manifesting as visible signs of aging (47).

Historically, the association of NRF2 activation with a protective response from damaging ROS has led to characterization of this response as a mode of cellular defense. Recently, however, NRF2 has also been demonstrated to play an important role in the wound healing process, extending our understanding of the importance of this process beyond defense to also include repair and healing of damaged skin (48–51). The importance of NRF2 in wound healing was first established in corneal and diabetic wounds, in which Nrf2 suppression resulted in delayed healing (49–51). More recently, Nrf2 dysfunction was further localized to the basal epidermis in diabetic wounds, and acute deletion of *Nrf2* in the basal epithelium was found to impair wound repair (48). Further mechanistic studies revealed Nrf2 activation induced Ccl2 expression to drive macrophage chemotaxis to the wound site to stimulate both the clean-up of cellular debris and induce keratinocyte proliferation via EGF release (48). Together, these findings suggest that epidermal Nrf2 is a primary regulator of the crosstalk events activated early after cutaneous injury, promoting the early-stage regenerative response in basal keratinocytes, ultimately stimulating re-epithelialization.

Through a temporal analysis of the effects of laser treatment on skin, we demonstrated NRF2 pathway activation in the early stages of the wound-repair response to fractional laser (Fraxel™) treatment, in which the NRF2 response is maximally activated at Day 1 and 3 post-treatment and declines to baseline over time (manuscript in preparation). This finding suggests that NRF2 may serve as a prominent conductor of repair not only in natural wound healing, but also in the context of a gold standard cosmetic rejuvenation procedure which inflicts damage activating the innate repair response of skin. We furthermore identified a synergistic activation of the N RF2 pathway by a combination of peptides + Nam in epidermal keratinocytes, in which the NRF2-mediated oxidative stress response was found to be amongst the top 10 most strongly activated synergy pathways by the combination of Pal-KTTKS, Ac-PPYL, and Nam. And interestingly, synergistic NRF2 activation was found to be further augmented by the incorporation of two additional peptides (Pal-KT and RGS) or with four peptides (Pal-KTTKS + Ac-PPYL + Pal-KT + RGS) alone in the absence of Nam. It is therefore possible that these combinations of materials may act to synergistically “prime” the regenerative capacity of skin cells, initiating a key NRF2/CCL2/EGF signaling axis involved in orchestrating tissue repair.

In addition to NRF2 activity, cellular regeneration requires large amounts of energy to fuel the repair and rebuilding processes, including the ATP required for proteasome and autophagy recycling processes, and the ATP expended in transcribing and translating genes into new proteins and associated export processes (52, 53). Our finding that the combination of peptides Pal-KT and Ac-PPYL synergistically restore in vitro cellular ATP levels that have been depleted due to the presence of ROS further highlights the beneficial cellular regeneration spaces in which peptide combinations operate.

It has been previously shown that peptide fragments result from damage to the ECM and thus are a “sensing” medium of extracellular matrix damage (17, 54). Cosmetically, peptides have been utilized in skin repair regimens to exploit this sensing mechanism of repair (55). However, the synergistic potential of peptides with diverse repair functions across the epidermis and dermis is not well understood. Our observation of the in vitro synergistic activation of NRF2 and energy-related processes by peptide combinations sheds more light on the specific mechanisms involved in damage and stress sensing to facilitate skin cell regeneration. Together, these findings provide additional insight into how specific combinations of bioactive peptides could be working, specifically, through “priming” the regenerative capacity of cells and facilitating the energetic requirements for cellular regeneration. As such, these peptide combinations could potentially mimic the beneficial effects of cosmetic rejuvenation approaches, without having to inflict additional damage.

---

**Supplemental Table 1:** qPCR Validated Synergies for combination of Nam, Pal-KTTKS, and Ac-PPYL.

| Nam (µg/mL) | Pal-KTTKS (µg/mL) | Ac-PPYL (µg/mL) | Gene | Observed Fold Change | Expected Fold Change | P-value |
| --- | --- | --- | --- | --- | --- | --- |
| 500 | 1 | 1 | GDE1 | 1.40 | 0.987 | 0.0198 |
| 500 | 1 | 1 | HMGCL | 1.59 | 1.08 | 0.0368 |
| 500 | 1 | 1 | MINPP1 | 1.38 | 1.02 | 0.0195 |
| 500 | 1 | 1 | SLFN5 | 1.77 | 1.09 | 0.0011 |

**Supplemental Table 2:** qPCR Validated Synergies for combination of Pal-KT & Ac-PPYL.

| Pal-KT (µg/mL) | Ac-PPYL (µg/mL) | Gene | Observed Fold Change | Expected Fold Change | P-value |
| --- | --- | --- | --- | --- | --- |
| 15 | 40.5 | MSMO1 | 9.62 | 3.15 | 4.46E-14 |
| 15 | 1 | PPARA | 2.54 | 1.09 | 1.23E-05 |

